# Kinesin-8 motors dimerize by folding their proximal tail domain into a compact helical bundle

**DOI:** 10.1101/2024.09.05.611543

**Authors:** Daria Trofimova, Caitlin Doubleday, Byron Hunter, Jesus Serrano Arevalo, Emma Davison, Eric Wen, Kim Munro, John S. Allingham

## Abstract

Kinesin-8 motor proteins help align and segregate chromosomes during mitosis by regulating the dynamics of kinetochore-attached microtubules and the length and position of the mitotic spindle. Some kinesin-8 isoforms accomplish these roles by operating as multifunctional mechanoenzymes that can traverse microtubules, accumulate at the microtubule plus-ends, and then remove terminal αβ-tubulin subunits. While these activities are mainly powered by the motor domain, whose unique structure-function relationships have been recently reported, the non-motor tail domain contains integral functional elements that have not been structurally illuminated. Using the *Candida albicans* Kip3 protein as a kinesin-8 model system, we present an X-ray crystal structure and hydrodynamic data showing how the motor domain-proximal segment of the tail directs the assembly of two kinesin-8 polypeptides into a homodimer that forms the stalk of this motor. Unlike the extended coiled coil-forming helices of most other motile kinesin stalks, the proximal tail of *Ca*Kip3 folds into a compact 92 Å-long four-helix bundle that dimerizes. The first and third helices provide most of the surface area for the dimer interface, while the other two helices brace the folded stalk structure. The upper and lower lobules of the helical bundle are separated by a flexible hinge that gives the exterior faces of the stalk slightly different shapes when bent. We propose that these unique characteristics provide structural rigidity to the kinesin-8 stalk, as well as sites for transient interactions with kinesin-8-associated proteins or other regulatory regions of the motor.

## Introduction

Cell division, intracellular transport, and muscle contraction are among the many biological activities that depend on kinesin, dynein, and myosin motor proteins. Although these enzymes use different mechanisms to move along their cytoskeletal trackways (kinesin and dynein move along microtubules; myosin moves along actin), the salient features they share include: (1) a catalytic motor domain that couples ATP binding, hydrolysis, and release of hydrolysis products to protein conformation changes, (2) a neck or neck linker domain that amplifies the motor domain-directed conformational changes to generate movement, and (3) a tail domain that mediates cargo or cytoskeleton binding^1^.

Motor protein oligomerization via the tail region is another functionally significant feature of cytoskeletal motors. Muscle and some non-muscle myosins form dimers by intertwining the extended alpha-helical regions of their tail into a flexible coiled coil that can be up to 500 Å-long^2, 3^. This interaction projects the motor domains outward to engage with actin filaments for muscle contraction or hand-over-hand walking along actin fibres. Similarly, many kinesins form extended dimeric complexes connected by a long coiled coil ‘stalk’ that lies in between their motor domain and the cargo-binding region of their tail domain^4^. This architecture can help coordinate long processive runs of individual kinesins, or teams of kinesins, along microtubules^5, 6^. Finally, cytoplasmic and axonemal dyneins form dimeric complexes via regions in their tail that are decorated with accessory proteins^7^. These complexes regulate dynein motility and enable minus end-directed transport of cargos along microtubules^8–10^.

Although X-ray crystallographic and electron microscopic structures of myosin and dynein tail domains are the most abundant, the number of structures elucidated for kinesin tail domains is growing. Sablin *et al*. published the crystal structure of a truncated kinesin-14 (Ncd) homodimer over 25 years ago^11^, showing two motor domains connected by an 43-residue-long coiled coil of the motor domain-proximal segment of the tail. The crystal structure of the hetero-tetramer-forming kinesin-1 motor (KIF5) determined by Kozielski *et al*. showed two motor domains connected via a 31-residue-long coiled coil interaction of their α-helical necks^12^. The crystal structure of the BASS domain of kinesin-5 (KLP61F, Eg5) solved by Nithianantham *et al.*^13^ showed how this kinesin forms homotetramers via an extended four-stranded coiled coil whose interface is ∼170 residues-long. More recently, the structures of a full-length autoinhibited kinesin-1 homodimer and a kinesin light chain-bound kinesin-1 heterotetramer were elucidated by Tan *et al*. using a combination of electron microscopy and AlphaFold structure prediction^14^. With further advances in protein structure prediction programs like DeepMind’s AlphaFold3^15^, these tools will give increasingly accurate glimpses of functional kinesin tail elements^14, 16^. This structural information is essential for understanding the regulatory mechanisms, cellular roles, and disease implications of these cytoskeletal motor proteins.

Compared to other kinesins, very little is known about the tail architecture of the kinesin-8 family of motors. These kinesins dimerize to provide transport and microtubule-length regulating functions in eukaryotes by using a form of bimodal operation that entails switching between processive motility along the microtubule lattice shaft and microtubule lattice disruption at the microtubule plus end^17–21^. This mechanistic difference from purely motile kinesins may be, in part, achieved through a unique mode of tail domain dimerization or tail structure dynamics, but no kinesin-8 tail structures have been determined experimentally. Thus far, it has been shown that the tail domain of *Saccharomyces cerevisiae* kinesin-8, *Sc*Kip3, has two discrete regions, dubbed the proximal and distal tail, that differentially control where and when *Sc*Kip3 functions in mitosis^20^.The proximal tail limits the microtubule-depolymerization activity of *Sc*Kip3 to astral microtubules that enter the vicinity of the bud neck, which helps cells reorient mispositioned anaphase spindles^22, 23^. The distal tail keeps *Sc*Kip3 from depolymerizing microtubules at the spindle midzone during anaphase, which prevents premature spindle disassembly until chromosome segregation is complete^22^. The molecular basis for these effects of the proximal and distal tail on *Sc*Kip3 activity are not yet known, nor are the structures of either domain.

In this study, we combined AlphaFold3 structure prediction, X-ray crystallography, analytical ultracentrifugation (AUC), and size-exclusion chromatography coupled to multiangle light scattering (SEC-MALS) to probe the structure and function of the motor domain-proximal region of the tail domain of *Candida albicans* Kip3, a close relative of *Sc*Kip3^24^. Our data show that *Ca*Kip3’s proximal tail forms a compact four-helix bundle that dimerizes in parallel with the same region of another *Ca*Kip3 subunit, producing a unique and compact kinesin stalk structure. The homologous region of *Sc*Kip3, *Schizosaccharomyces pombe* Klp5/Klp6, *Drosophila melanogaster* Klp67A, *Homo sapiens* Kif18A, and *Homo sapiens* Kif19 form similar stalk structures according to AlphaFold3 predictions. Although the alpha-helical region preceding the proximal tail bundle bears some coiled coil-forming sequence characteristics, our data suggest that this region is not a significant contributor to motor dimerization, unlike in kinesins-1, 2, 5, and 7^25^.

## Results

### The proximal tail forms the dimerization domain of kinesin-8

Kinesin tails usually contain amino acid sequence patterns that allow them to form dimers via intermolecular coiled coils (**Supplementary Figure 1A-1C**)^25^. These regions are distinguished by a repeating series of seven-residue blocks (heptads) that contain nonpolar amino acids (often Leucine or Isoleucine) in position 1 and 4; and charged amino acids at position 5 and 7^26^. For metazoan and fungal kinesin-8s, the heptad signature of the tail region is imperfect and is confined to a short, 30 to 40-residue segment of the tail near the motor domain, raising questions about how kinesin-8s form functional motor dimers (**Supplementary Figure 1D**). AlphaFold3 predicts that this region, which we named the neck extension (NE), does not intertwine into a canonical coiled coil with its partner subunit. Instead, it continues into a compact four-helix bundle that embraces its partner subunit in parallel, forming a proximal tail (PT) homodimer (**Figure 1**). The remaining C-terminus of the protein, which forms the distal tail (DT), is mostly unstructured (**Figure 2A**).

**Figure 1.**
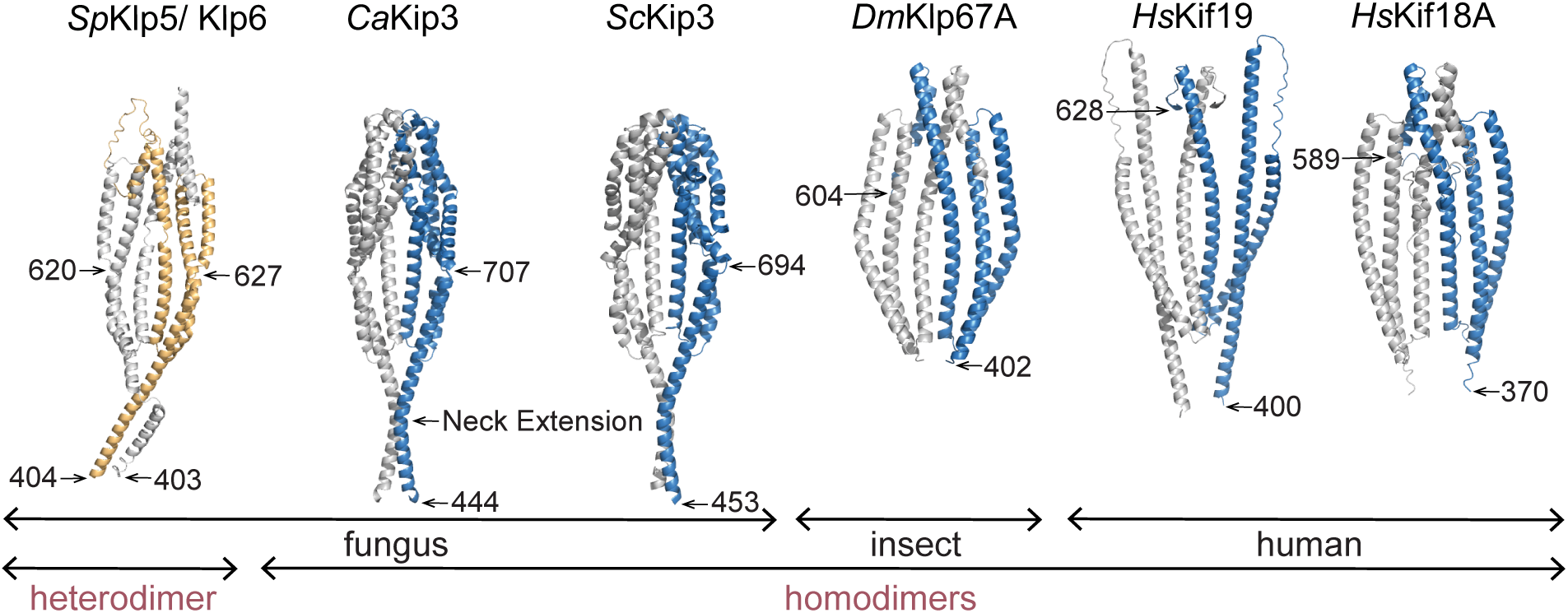
Dimer structures of proximal tail of kinesin-8s predicted by AlphaFold3. AlphaFold3 structural models of *Saccharomyces cerevisiae* Kip3, *Candida albicans* Kip3, *Homo sapiens* Kif18A, *Schizosaccharomyces pombe* Klp5/Klp6 heterodimer, *Drosophila melanogaster* Klp67A, and *Homo sapiens* Kif19. In the model of *Sp*Klp5/Klp6 heterodimer Klp5 is colored in yellow, Klp6 is grey. For all other models, one protein chain is shown in blue, and the other is in light grey. The motor domains and low confidence loops were removed.

**Figure 2.**
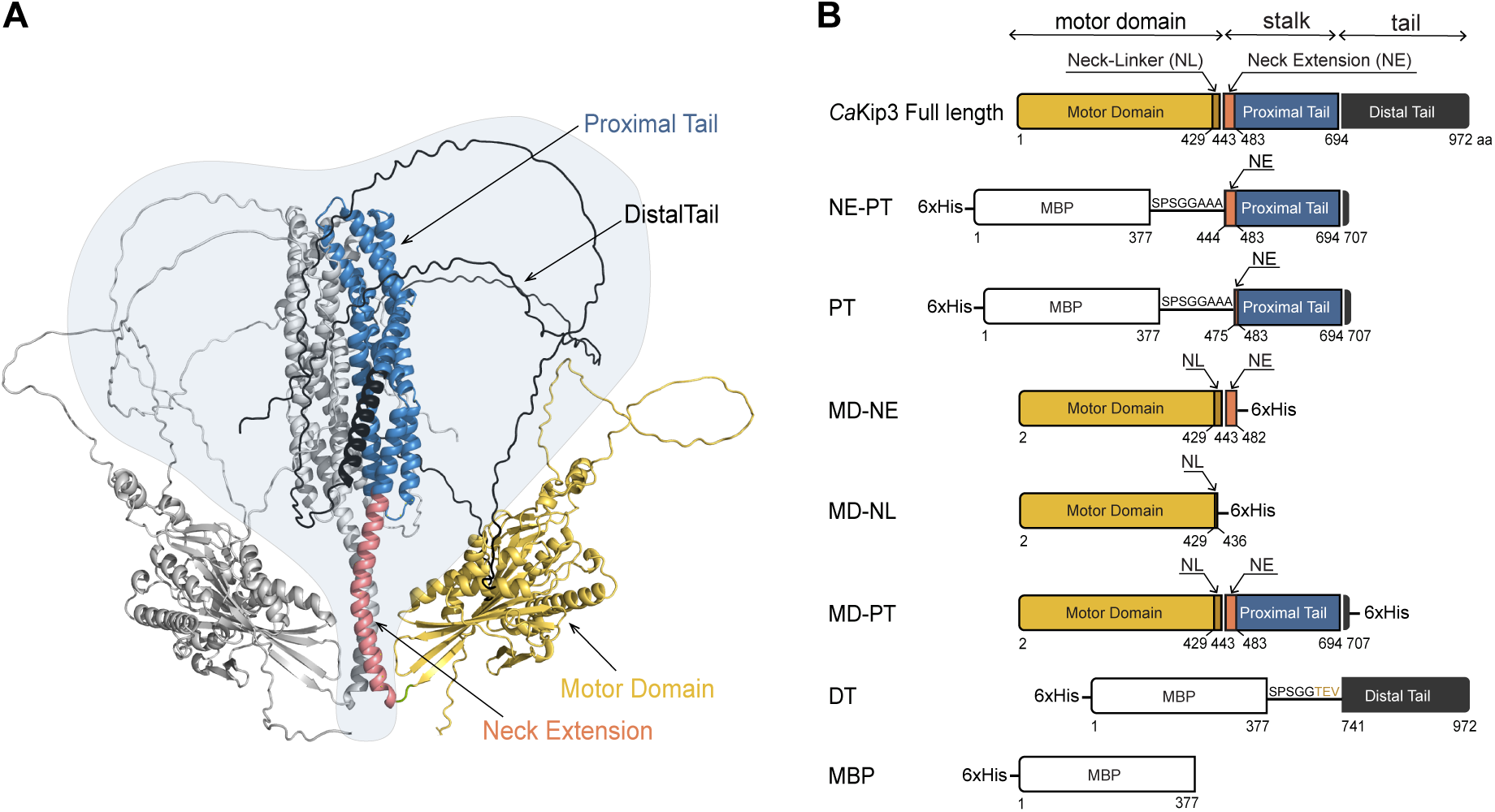
AlphaFold3 model of the entire *Ca*Kip3 dimer and diagrams of truncated *Ca*Kip3 constructs. (A) Predicted structure of a full *Ca*Kip3 dimer. One monomer is colored in light grey; the second monomer is colored in accordance with the domain map in (B). A light blue blob is included to highlight the unstructured tail domain. (B) Domain boundary diagram of *Ca*Kip3 showing the motor domain (MD), neck linker (NL), neck extension (NE), proximal tail domain (PT), and distal tail domain (DT). The components of the protein constructs used for the structural and/or functional studies are shown below.

To experimentally test AlphaFold3’s predictions, we cloned, expressed, and purified six fragments of *Candida albicans* kinesin-8 (*Ca*Kip3) (**Figure 2B**) using BL21 *E. coli* cells as an expression system. The boundaries of these constructs were selected based on the AlphaFold3 model of the *Ca*Kip3 dimer. The *Ca*Kip3 MD-PT construct includes the motor domain (MD), neck linker (NL), neck extension (NE), and the proximal tail domain (PT) (residues 1-707). *Ca*Kip3 MD-NE includes the motor domain, neck linker, and neck extension (residues 1–482). *Ca*Kip3 MD-NL contains the motor domain and a slightly truncated neck linker (residues 1–436). These three constructs also include a C-terminal 6x His-tag for Ni-NTA agarose affinity chromatography purification. The *Ca*Kip3 NE-PT construct contains the neck extension and proximal tail (residues 444-707). The *Ca*Kip3 PT construct contains the proximal tail only (residues 475-707), and *Ca*Kip3 DT contains the distal tail only (residues 741-972). These latter three constructs also include a 6x His tag and maltose-binding protein (MBP) fused to their N-terminus for Ni-NTA agarose affinity purification and to improve their solubility, respectively. SDS-PAGE analysis of the purified proteins is shown in **Supplementary Figure 2A**.

The oligomeric composition of these purified proteins was determined by light scattering (SEC-MALS) and by sedimentation analysis (AUC). Both analyses showed that most of the molecules contained within the purified *Ca*Kip3 MD-NE and MD-NL constructs have masses that closely match the theoretical molecular weight of a protein monomer (**Figure 3A, 3B, 3G and Supplementary Figure 2B, 2C**). The same was observed for a construct that contains MBP only (**Figure 2B, 3F, 3G and Supplementary Figure 2F**). In contrast, the observed masses of the molecules in the *Ca*Kip3 NE-PT, PT, and MD-PT-containing samples was roughly double the theoretical molecular weight of these constructs, confirming that proximal tail region forms homodimers (**Figure 3C, 3D, 3E, 3G and Supplementary Figure 2D, 2E**). These data also show that the neck extension sequence that contains the predicted coiled coil-forming domain (residues 444-471) was not necessary to produce dimers of the proximal tail (*Ca*Kip3 PT construct) and that its inclusion did not produce dimers of the motor domain construct (*Ca*Kip3 MD-NE). Rather, only with inclusion of the proximal tail (*Ca*Kip3 MD-PT) did we observe motor domain dimers (**Figure 3E**). These data demonstrate that the proximal tail domain alone forms the main interface for *Ca*Kip3 dimerization.

**Figure 3.**
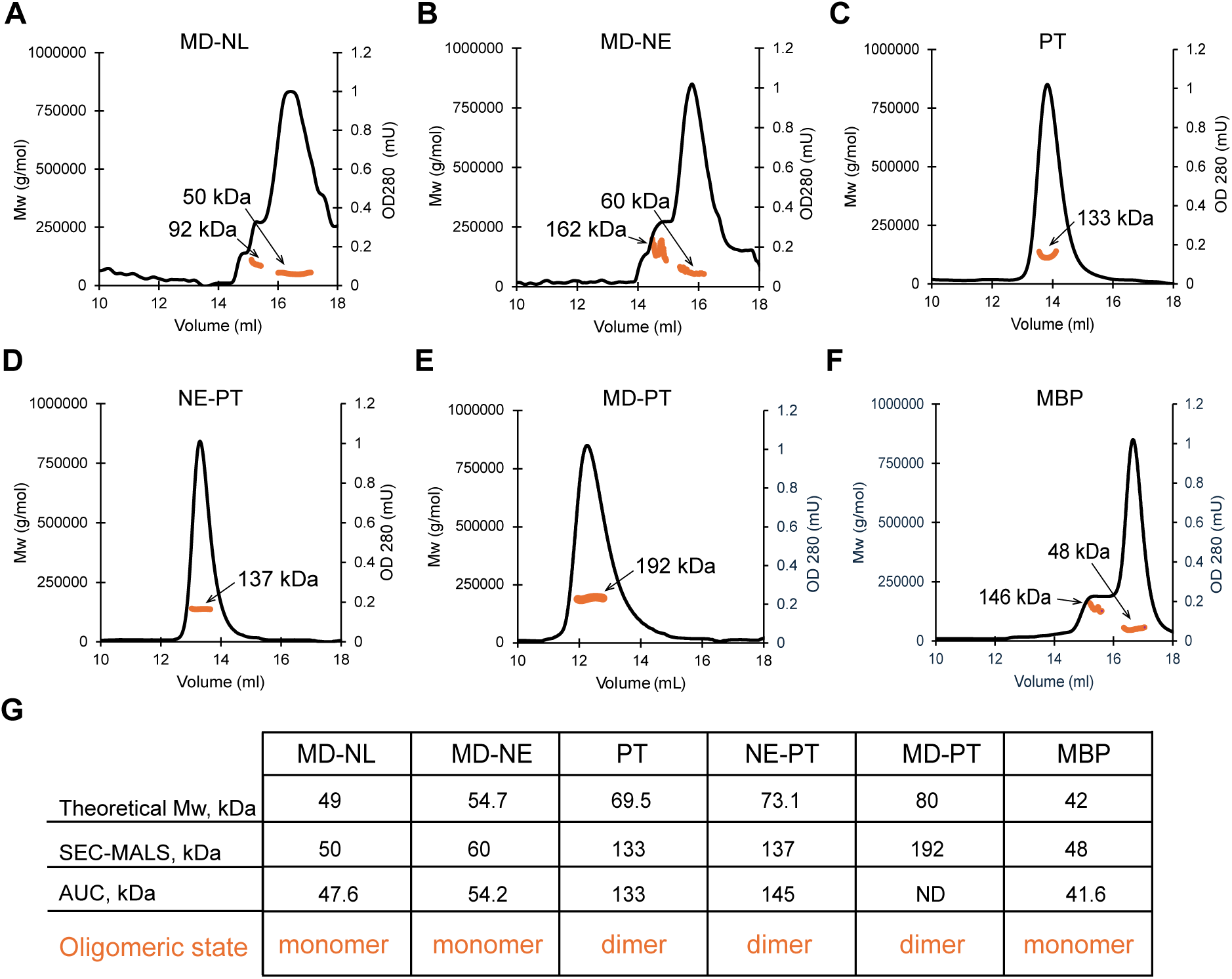
SEC-MALS analysis of the oligomeric state of *Ca*Kip3 constructs. (A-F) Size Exclusion Chromatography – Multi-Angle Light Scattering (SEC-MALS) analysis of 20 μM solutions of (A) Motor Domain-Neck Linker (MD-NL), (B) Motor Domain-Neck Extension (MD-NE), (C) Proximal Tail domain (PT), (D) Neck Extension-Proximal Tail domain (NE-PT), (E) Motor Domain Proximal Tail (MD-PT), and (E) MBP proteins in HEPES 20 mM, MgCl_2_ 1 mM, NaCl 150 mM, DTT 1 mM, pH 7.0 buffer. (G) Summary of theoretical and experimentally determined Mw of all constructs with SEC-MALS, and analytical ultracentrifugation (AUC) techniques.

### Molecular architecture of the CaKip3 stalk

To acquire an experimentally derived structure the *Ca*Kip3 proximal tail, we subjected the MBP-tagged *Ca*Kip3 NE-PT and *Ca*Kip3 PT constructs to sparce-matrix crystallization screening by sitting-drop vapour diffusion. Crystals grew for both constructs within one week, but only the *Ca*Kip3 PT crystals diffracted X-rays sufficiently for structure determination. Synchrotron diffraction data was collected on single *Ca*Kip3-PT crystals and the structure of the protein was solved to a resolution of 2.5 Å by automated molecular replacement (MR) using one subunit of the AlphaFold3-predicted *Ca*Kip3 PT dimer as a search model. The top MR solution comprised eight copies of the *Ca*Kip3 PT subunit in the asymmetric unit (subunits A-H). Each subunit consists of a four-helix bundle that closely matches the AlphaFold3 model but bears no structural similarity to any protein structure in Protein Data Bank (PDB) (**Figure 4A**). Like the AlphaFold3 dimer model, two *Ca*Kip3 PT subunits embrace each other in parallel via their first and third alpha-helices, so that the N termini extend outward in the same direction (**Figure 4B, 4C**). This configuration would hold the two motor domains in proximity. There are four of these *Ca*Kip3 PT dimers in the crystallographic asymmetric unit (dimers AB, CD, EF, and GH) (**Figure 4D**). Dimers AB and CD associate in an antiparallel orientation in one plane, and dimers EF and GH are similarly arranged but lay on a tangential plane angled about 60° from the AB and CD dimers.

**Figure 4.**
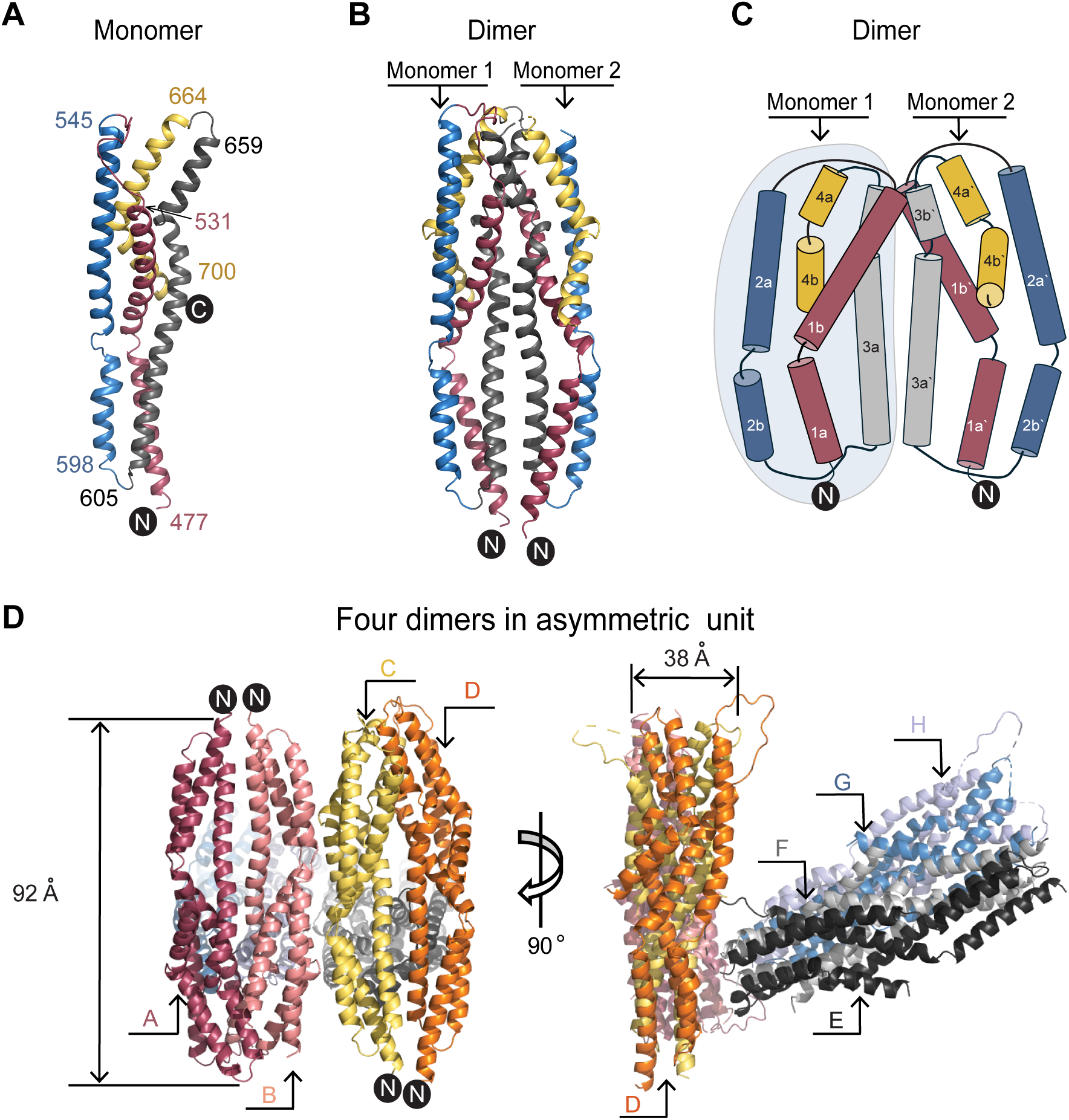
Crystal structure of the *Ca*Kip3 proximal tail. (A) Cartoon representation of the monomer of the *Ca*Kip3 PT, where each individual alpha-helix is colored separately. Helix 1 (raspberry red) consists of residues 477 to 531, helix 2 (blue) consists of residues 545 to 598, helix 3 (grey) consists of residues 605 to 659, and helix 4 (yellow) consists of residues 664 to 700. The N and C -termini are labeled. (B) Cartoon representation of the dimer AB of the *Ca*Kip3 PT structure. (C) Cylinders representation of the dimer of the *Ca*Kip3 PT. Alpha-helices are represented as colored cylinders, and connecting loops are thin lines. Numbers with apostrophes correspond to the helices of a second monomer, B. The dimerization interface is formed via helices 1b, 3a, and 3b. (D) Orientation of the four dimers in asymmetric unit of the crystal. Dimer 1 consists of monomer A (raspberry red) and B (deep salmon red). Dimer 2 consists of monomer C (yellow) and D (orange). Dimer 3 consists of monomer E (black) and F (grey). Dimer 4 consists of monomer G (sky blue) and H (light blue). Cartoon representations were generated with PyMOL (DeLano Scientific LLC) ^55^.

Unexpectedly, there was no discernable density for the maltose-binding protein in the refined electron density map, nor was their volume to accommodate it among the *Ca*Kip3 PT molecules in the crystallographic lattice. To account for the absence of MBP, we examined the protein content of the crystals by SDS-PAGE. Although we collected dozens of crystals and rigorously removed all precipitant solution from them prior to dissolving them and denaturing their contents in SDS-PAGE running buffer, we were unable to resolve specific bands of proteins on the gels (**Supplementary Figure 3A**). However, we found that incubating the protein at room temperature in buffer or crystallization solution for several days produced at least five prominent proteolytic species (52, 42, 27, and 18 kDa) that could be resolved by SDS-PAGE after Coomassie staining (**Supplementary Figure 3C**). Western blotting analysis with an anti-His antibody detected the 69 kDa, 52 kDa, and 42 kDa bands, but did not detect the 27 kDa and 18 kDa bands (**Supplementary Figure 3D**). Based on these observations, we predict that cleavage of the *Ca*Kip3 PT construct took place at the N-terminus of the proximal tail domain and somewhere in the N-terminal half of the proximal tail. This would give 42 and 52 kDa fragments containing the His-tag and MBP domain that were detected by Western blotting. The 27 and 18 kDa fragments presumably contain the proximal tail but no His tag for detection by Western blotting (**Supplementary Figure 3B**). Moreover, the size of the proximal tail protein we were able to model in the crystal lattice is approximately 27 kDa. These analyses provide evidence that MBP-tagged *Ca*Kip3 PT fusion protein had been proteolyzed prior to crystallization, separating the proximal tail module from segments containing the His-tagged MBP component.

### The dimerization interface

According to PDBePISA, an interactive tool for the exploration of macromolecular interfaces^27^, the interaction between two proximal tail subunits buries up to 2,817 Å^2^ of the 15,216.8 Å^2^ total surface area of each subunit. A plot of the electrostatic surface potential of the *Ca*Kip3 PT monomer reveals two stretches of non-polar surface spanning the vertical axis (**Figure 5A, 5B**). These two areas are situated on opposite sides of the monomer, one at the dimerization surface and the other on the interdimer contact site in the crystal, indicating that the *Ca*Kip3 PT dimer complex is stabilized by the hydrophobic effect. At an X-ray diffraction data resolution of 2.5 Å, the electron density maps were sufficiently detailed to model most of the proximal tail residue side chains that form the bonding interactions to stabilize the *Ca*Kip3-PT dimer (**Figure 5C**). Using the Analysis by Protein Contacts Atlas tool, we identified ∼102 contacts between the subunits^28^. Helices 1b, 3a, and 3b provide most of the contacts that hold the proximal tail dimer together. Helix 3a interacts with its counterpart in the other monomer (helix 3a’) and with residues in helix 1b’. The interactions are formed by both polar and non-polar residues, some of which are conserved in the *S. cerevisiae* and *S. pombe* kinesin-8s (green and blue dots in **Supplementary Figure 4**). In the *Ca*Kip3 PT, six residues form salt bridges between these helices (**Figure 5D**). These involve Arg 515 and Asp 635, Lys 516 and Asp 635, and Arg 620 and Glu 621. Notably, the AlphaFold3 predicted structure of the *Sc*Kip3 PT dimer has polar groups at similar locations on helices 1b and 3a (**Supplementary Figure 4 and 5A**). In the predicted *Sc*Kip3 PT dimer model, Asn 628 bonds with Asn 628 of another monomer in the middle of helices 3a and 3a’, and Arg 527and Glu 639 mimic the upper interactions between helix 3a and 1b’ seen in the *Ca*Kip3 PT structure. The AlphaFold-predicted *Sp*Klp5/6 PT heterodimer model also has polar groups at those sites on helices 1b’ and 3a (**Supplementary Figure 5B**). Here, Asn 576 and Glu 575 of Klp5 interact with Glu 569 and Asn 570 of Klp6 in the middle of helices 3a and 3a’, and Asn 583 of Klp5 and Lys 576 of Klp5 interact between helix 3a and 1b’. Interestingly, *Sp*Klp5 E575P and *Sp*Klp6 E569P mutants of this kinesin-8 are non-functional, and cause an increase in pre-anaphase spindle length, duration, and severity of spindle protrusion length in *S. pombe* cells^29^. These observations suggest that polar and electrostatic interactions in the proximal tail complex are important for kinesin-8 function.

**Figure 5.**
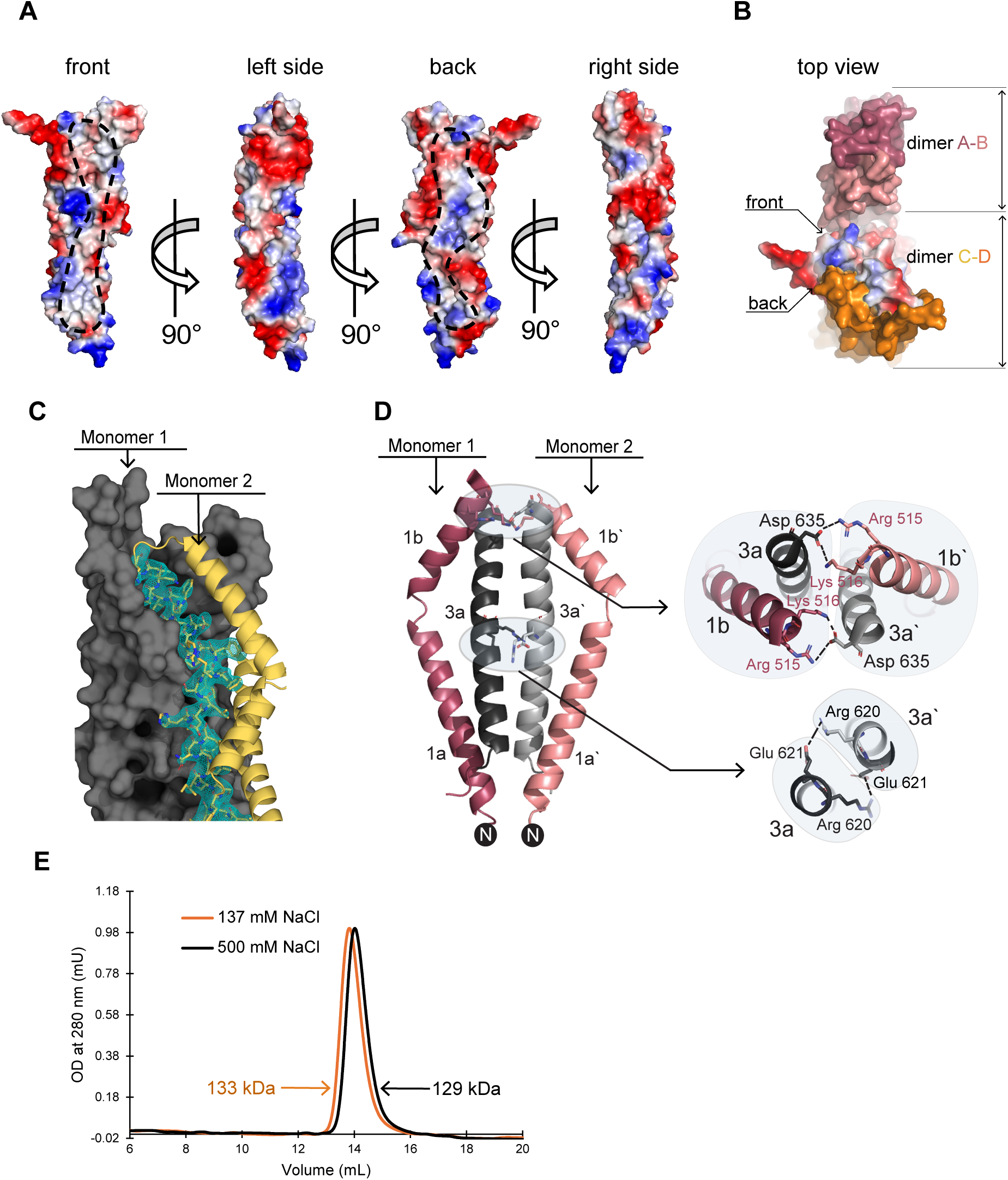
*Ca*Kip3 proximal tail dimer interface. (A) Surface charge representation of the *Ca*Kip3 PT with positively and negatively charged areas in blue and red, respectively. Electrostatic surface visualization was generated with PyMOL. Non-charged nonpolar area of the front and back surface of a monomer is circled with dashed lines. This is an area where the monomer of an intradimer binds (front) or another dimer (back) binds forming interdimer interface. (B) A top view of the interdimer and intradimer contacts observed in the *Ca*Kip3 PT crystal. One dimer is formed by chain A and B, another by C and D. (C) The representation of the *F*_obs_ − *F*_calc_ omit map (contoured at 3.0 *σ,* green-blue mesh) of helix 3a calculated after deletion of the helix from final model. Monomer 1 shown as grey surface, Monomer 2 as yellow cartoon. (D) Cartoon representation of dimerization interface organized mainly by 3a helices of monomer A and B. Helices 1b interact with 3a at the top site of the bundles. Salt bridges are shown as dashed lines. Grey half circles on the zoon in views represent individual monomers in the structure of the dimer. (E) SEC-MALS profiles of *Ca*Kip3 PT at low (137 mM NaCl – orange line) and high (500 mM -black line) ionic strength.

To investigate the importance of the electrostatic interactions in *Ca*Kip3-PT dimerization, we performed SEC-MALS analysis on the *Ca*Kip3 PT construct in low and high ionic strength buffers. The SEC chromatographs of *Ca*Kip3 PT dimers show single peaks with very similar elution profiles when the buffer conditions include 137 mM NaCl (orange line) and 500 mM NaCl (black line) (**Figure 5E**). MALS analysis of the molecules in these peak fractions gives molecular weights of 133 kDa (137 mM NaCl) and 129 kDa (500 mM NaCl), nearly equal to the theoretic mass of a *Ca*Kip3 PT dimer. This data indicates that electrostatic interactions at the dimer interface are not the main contributors to dimerization.

### Intramolecular contacts in the folded CaKip3 proximal tail monomer

ProteinTools, a web server toolkit for protein structure analysis^30^, indicates that the four-helix bundle of *Ca*Kip3 PT monomers is stabilized by three salt bridges and numerous hydrophobic interactions between helices (**Figure 6A, 6B**). There are two salt bridges in upper lobule of *Ca*Kip3 PT and one in the lower lobule. The interaction between Arg 532 and Glu 559 in *Ca*Kip3 is similarly positioned to the Arg 562 and Glu 545 salt bridge in the AlphaFold3 model of the *Sc*Kip3 PT (**Figure 6A**). The hydrophobic contacts distribute evenly along the lengths of helices 1, 2 and 3, and are distributed similarly in the AlphaFold3 model of the *Sc*Kip3 PT (**Figure 6B**). Non-polar residues that share similar positions in the lower lobule of the *Ca*Kip3 PT and *Sc*Kip3 PT bundle include Leu 582 (Leu 585 in *Sc*Kip3), Leu586 (Ile 589), Ile 615 (Leu 619), Leu 491 (Val 500), and Leu 487 (Ile 496). In the upper lobule, Leu 679 (Ile 669), Ile 551 (Ile 523), Leu 518 (Leu 530), Leu 572 (lle 575), Leu 569 (Ile 572), Ile 565 (Leu 568), Leu 561 (Ile 564), and Ile 554 (Val 557) share equivalent positions in *Ca*Kip3 and *Sc*Kip3.

**Figure 6.**
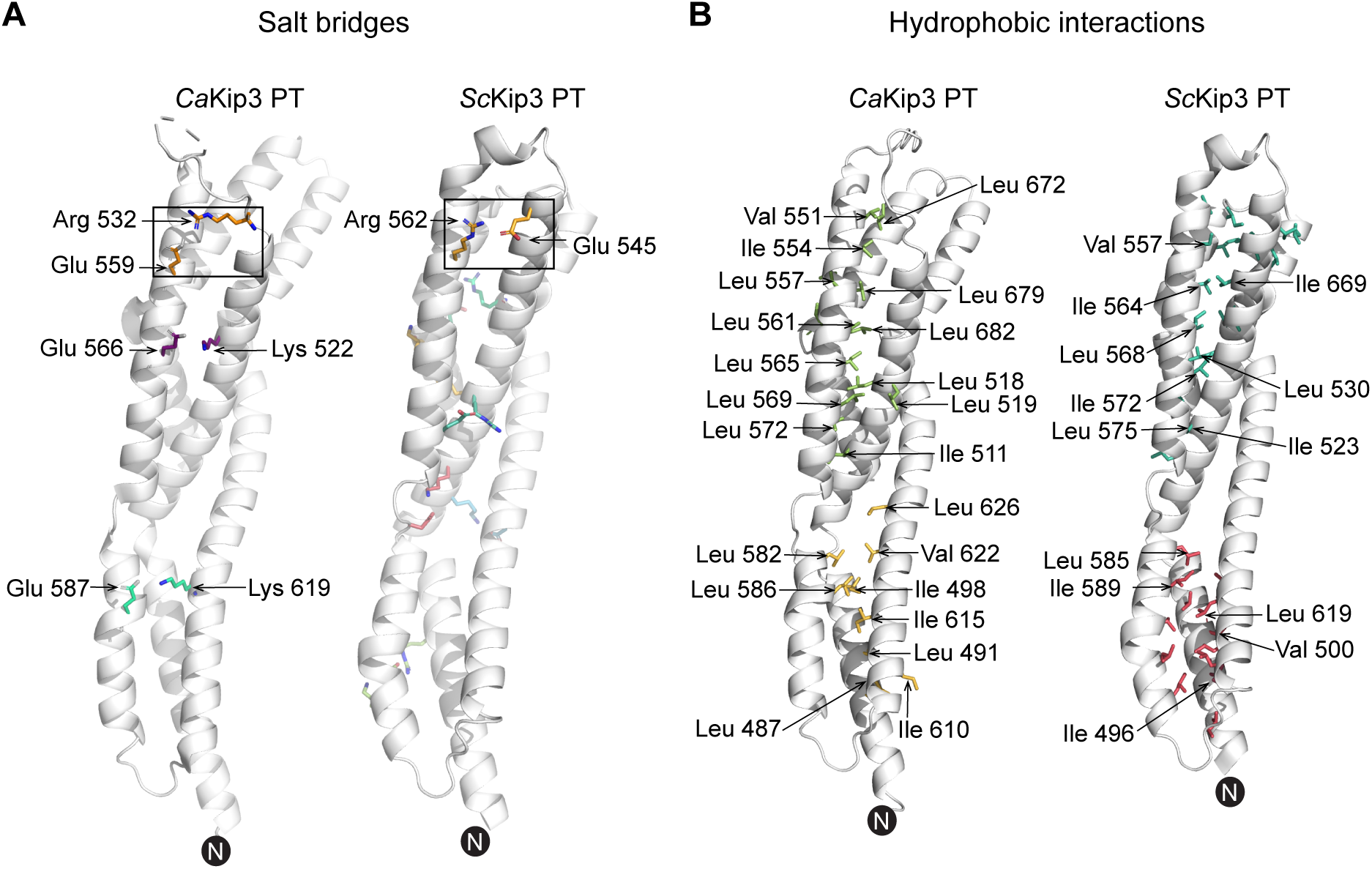
Intramolecular interactions that stabilize the *Ca*Kip3 PT monomer fold. (A) Salt bridges which stabilize monomers of *Ca*Kip3 and *Sc*Kip3 proximal tail domain fold are presented as coloured sticks and labelled with amino acid numbers. Salt bridges located at the same position are shown in the box. (B) Amino acids involved in hydrophobic interactions in monomers of *Ca*Kip3 and *Sc*Kip3 are shown as sticks. Lower and upper hydrophobic clusters are coloured in different colours. *Sc*Kip3 amino acids which situate at similar positions in the *Ca*Kip3 structure are labelled. The N terminus is marked. Identification of salt bridges and hydrophobic cluster were done with ProteinTools application^30^.

### The CaKip3 proximal tail can bend

When we were modeling the structures of each subunit (A-H) in the *Ca*Kip3 PT crystal, we noticed numerous differences in the conformation of the loops connecting the helices, as well as differences in the relative angles of some helices. Indeed, the RMSD values obtained by superimposing the C-alpha atoms of each subunit onto the other seven subunits range from 0.356-1.749 Å. While the flexible loops connecting the alpha-helices account for some of this structural deviation, we discovered that the conformation of each of the crystallographic dimers is slightly different (**Figure 7**). Moreover, the average B-factors for each chain differed as well, with chains G and H having the highest B-factors. These findings imply flexibility in the proximal tail fold. When we used the “Morph” tool in PyMOL to generate an interpolated trajectory of a *Ca*Kip3 PT subunit transitioning between the two most dissimilar conformations, we identified a “flexible” hinge between the upper and lower lobules of the proximal tail four-helix bundle (**Figure 7**). This hinge is formed by the segments that separate helices 1a and 1b, and helices 2a and 2b, and by a flexible sequence in the middle of helix 3a (yellow highlighted sequences in **Supplementary Figure 4**). Movement of the lower lobule relative to the upper lobule via this hinge leads to a displacement of the ends of helices 1a, 2b and 3a by 9.4 Å, 8.2 Å, and 7.3 Å, respectively. When the morph is applied to two *Ca*Kip3 PT subunits that are interacting as a dimer (A transitioning to B in one half of the dimer, and B transitioning to A in the other half, so they move complementarily to each other), the *Ca*Kip3 PT dimer bends enough to give the exterior faces of the homodimer slightly more concave or convex shapes.

**Figure 7.**
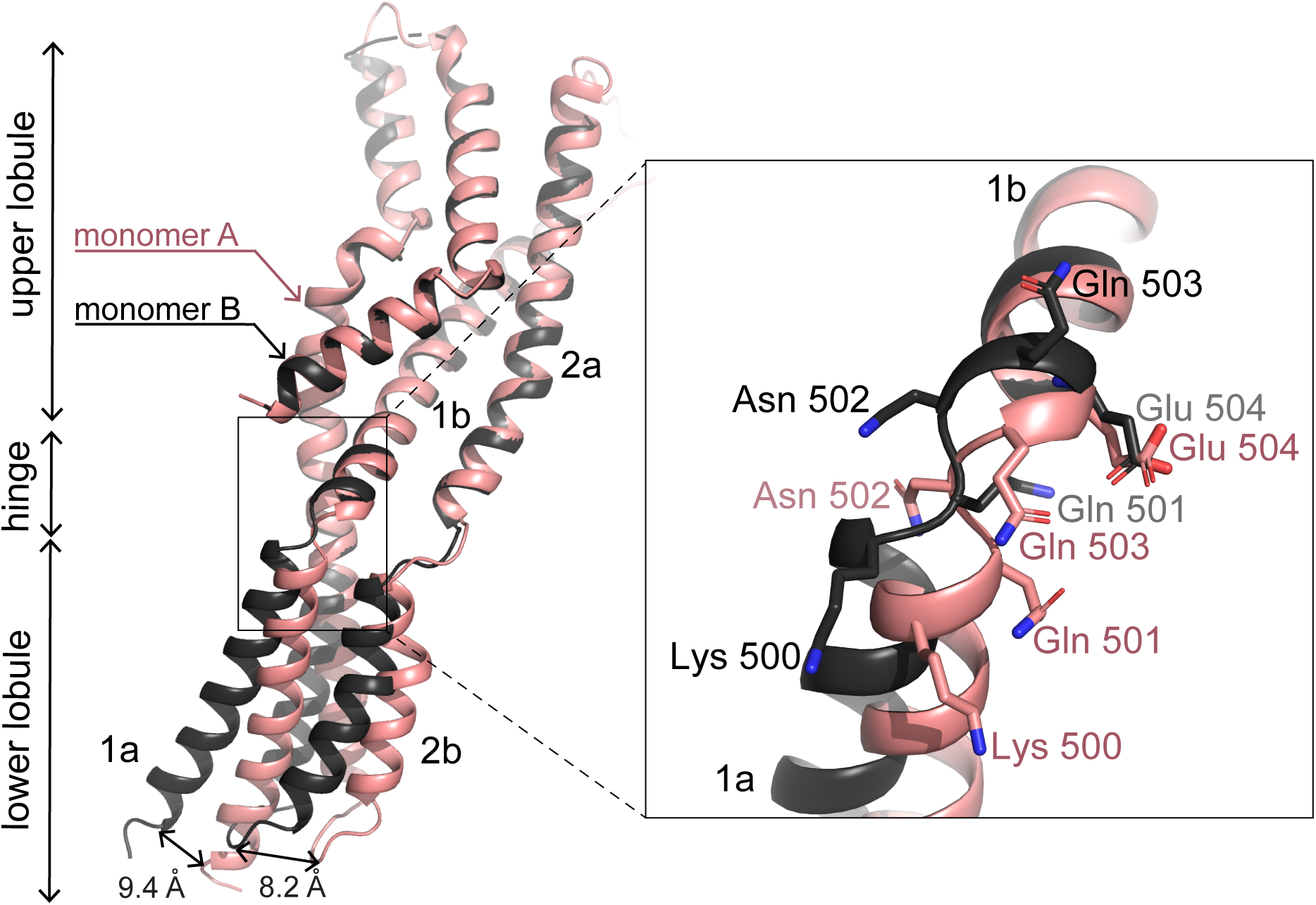
Bending of *Ca*Kip3 proximal tail subunits. Monomer A (deep salmon red) and B (black) were superimposed via their 1b helix. The displacement of the bottom part of helix 1a for monomers A and B was measured using the α-carbon of Asp 478 and the displacement of the bottom part of helix 2b for monomers A and B was measured using the α-carbon of Asp 599. The zoom-in the box shows the hinge between helix 1a and 1b where divergence in the location of helix 1a and 1b begins. Amino acids with the highest variation in orientation are shown in sticks representation.

### The proximal tail is not directly involved in microtubule interactions

The tail domain of some kinesin-8s can enhance processive motility, promote microtubule plus-end binding, and stabilize microtubules, indicating that the tail domain contains a microtubule binding site^31–33^. Recent studies have confirmed this for residues 802-898 of the tail of Kif18A^34^, and residues 481-805 of *Sc*Kip3^35^, but the exact microtubule binding surface within these regions are currently unknown. To determine whether the proximal tail of *Ca*Kip3 contains a microtubule binding interface, we performed a microtubule binding assay with the *Ca*Kip3 NE-PT and *Ca*Kip3 PT constructs, and with a construct that includes the distal tail of *Ca*Kip3 (*Ca*Kip3 DT; residues 740-972aa) (**Figure 2B and Figure 8**). In this assay, the *Ca*Kip3 proteins were incubated with increasing concentrations of taxol-stabilized microtubules and then centrifuged to pellet the microtubules and any kinesin protein that bound to them. SDS-PAGE analysis of the supernatant and pellet fractions was then used to identify the distribution of the kinesin protein in relation to the pelleted microtubules. We observed a small fraction of the input *Ca*Kip3 PT and *Ca*Kip3 NE-PT constructs in the pellet with the microtubules, however, the abundance of these proteins did not change with increasing concentration of microtubules (**Figure 8B, 8D**). This observation indicates that the *Ca*Kip3 PT and *Ca*Kip3 NE-PT proteins do not bind to microtubules specifically. In contrast, the abundance of *Ca*Kip3 DT protein in the pellet increased with the microtubule concentration (**Figure 8E, 8F**). These data show that *Ca*Kip3 proximal tail proteins with and without the first predicted coiled coil-forming domain do not contain a microtubule binding site, while the distal tail does.

**Figure 8.**
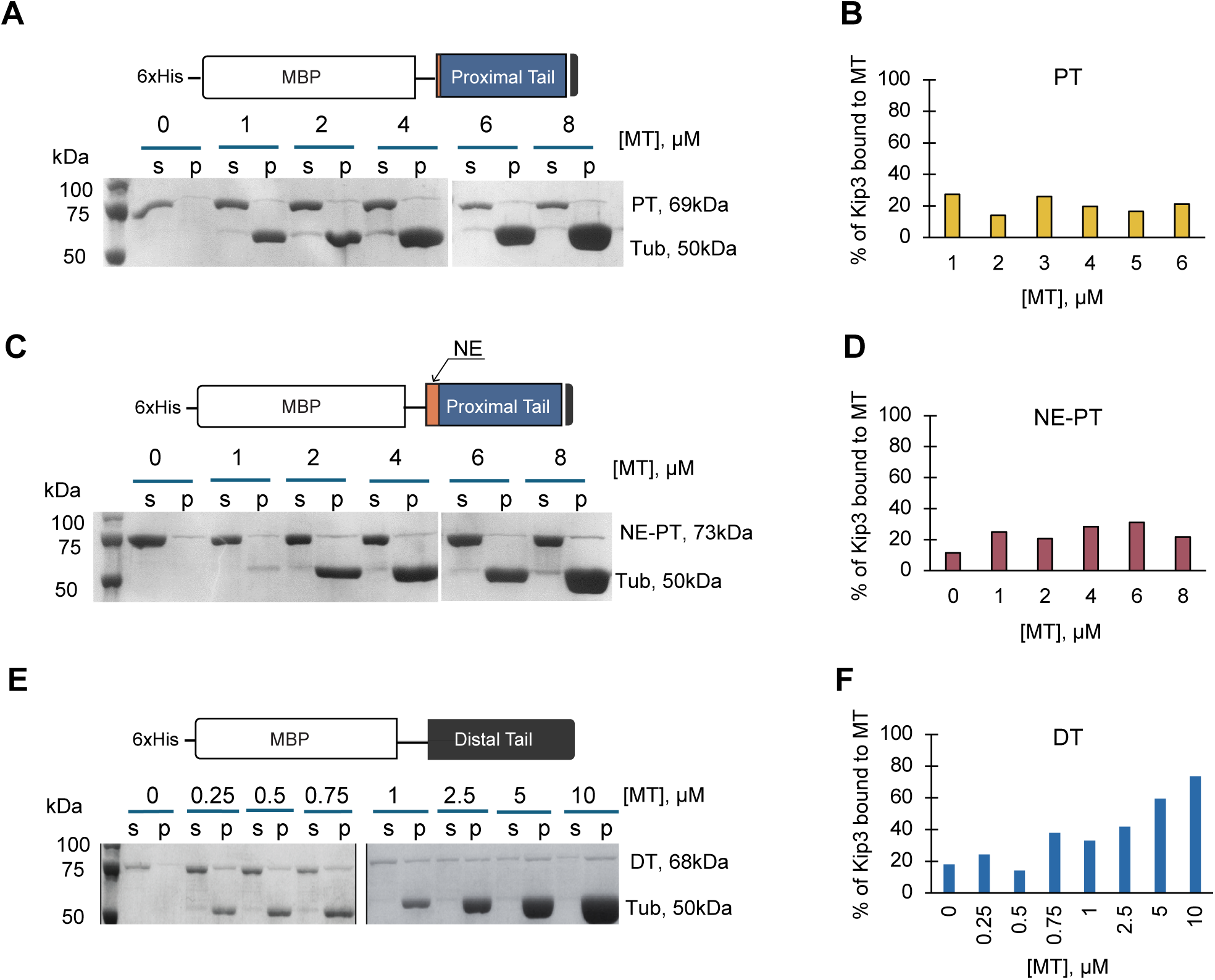
Microtubule binding properties of the proximal and distal tail. Representative SDS-PAGE gels of the microtubule binding assay for the *Ca*Kip3 PT (A), Neck-Extension Proximal Tail (NE-PT) (C), and Distal Tail (DT) (E) constructs. Kinesin proteins at fixed concentrations of 1 μM were incubated in the presence of 2 mM AMP-PNP and increasing concentrations (0-10 μM) of taxol-stabilized microtubules (calculated based on tubulin dimer concentration) in BRB80 buffer supplemented with 300 mM KCl for 20 min. The binding reactions were pelleted by centrifugation to separate the free kinesin (S) and microtubule-bound kinesin (P). SDS-PAGE gels stained with Coomassie brilliant blue were used to determine the fraction of microtubule-bound protein with the help of ImageJ software (B, D, and F). The percentage of bound kinesin was plotted as a function of microtubule concentration.

## Discussion

This study of the structure and function of the tail region of the *Candida albicans* kinesin-8, *Ca*Kip3, revealed that the proximal tail domain folds into a compact helical bundle that dimerizes, forming a broad interface (2,817 Å^2^) between two *Ca*Kip3 motor subunits. Although the adjacent neck extension domain contains sequence features of a coiled coil-forming alpha-helix, this region of *Ca*Kip3 does not appear to contribute to the dimer’s assembly or stability, according to our hydrodynamic analyses of the constructs that include this region. These findings support the AlphaFold3 predictions for the *Ca*Kip3 dimer structure. They also suggest that the connection between the motor domain and dimerization interface of kinesin-8s is separated by dozens of residues. Based on the AlphaFold3 predictions of other kinesin-8s, these are hallmark features of this family of motors.

We propose that the stalks of kinesin-8s have evolved to have a larger dimerization interface closer to their C-terminus, and a weaker, more flexible coiled coil near their motor domain compared to other kinesins. These characteristics could enable the bimodal mechanism (motility and microtubule depolymerization) of kinesin-8s in several ways. First, a more flexible stalk near the motor domain would benefit protofilament switching events that kinesin-8 uses to avoid microtubule bound obstacles on its way to the plus end^36–38^. In line with this thinking, the extended neck-linker of *Hs*Kif18A greatly enhances the ability of the motor to sidestep obstacles on microtubules to maintain processive movement^38^. Second, added flexibility in the kinesin-8 stalk near the motor domain may enable both motor domains of a dimer to adopt the nucleotide pocket closed, pro-depolymerization state on curved protofilaments at microtubule ends, assuming that dimeric kinesin-8s arrange their two motor domains in a head-to-tail fashion at the microtubule end^24^. Experiments suggest that this should not be possible for motile kinesins because the neck-linker of the forward head would be pulled backwards, preventing forward head nucleotide pocket closure^39^. However, if the proximal tail dimerization interface provides additional structural flexibility to the stalk, it may be possible for a dimeric kinesin-8 to have both motor domains on the same protofilament with closed nucleotide pockets and docked neck-linkers, in contrast to motile motors. For *Ca*Kip3 specifically, this would allow loop-1 contacts to form between and within dimers^24^.

The presence of multiple copies of the *Ca*Kip3 PT molecule in the asymmetric unit of the crystal was advantageous because it showed that the rigid upper and lower lobules of the helical bundle are separated by a flexible hinge. Although we do not yet understand if or how conformational transitions in the proximal tail domain are involved in kinesin-8 motility or microtubule depolymerization, one possibility is that they may facilitate the transduction of conformational changes between the catalytic motor domains. Another possibility is that the bendability of the proximal tail provides dynamic sites for transient interactions with kinesin-8-associated proteins or other regulatory regions of the motor. Indeed, recent studies have shown that these regions provide more than just an interaction interface for the catalytic motor subunits^40^. The non-motor tail of kinesins^41, 42^, myosins^43^ and dyneins^44^ can regulate the activity of the motor by folding into a variety of compacted structures that are stabilized by intra or intermolecular coiled coils. Upon modification with post-translational modifications and/or regulatory protein interactions, these structures rearrange to form extended tails, leading to motor activation^45–47^. Further studies that focus on identifying regulatory factors of *Ca*Kip3 and examining the motility and microtubule remodelling activities of dimeric *Ca*Kip3 motors with targeted mutations in the proximal tail domain will be needed to probe these possibilities.

## Materials and Methods

### Sequence analysis and Structure prediction

Protein sequences for the *Ca*Kip3, *Sc*Kip3, *Sp*Klp5/6, *Dm*Klp67A, *Hs*Kif18A, and *Hs*Kif19 were retrieved from UniProt^48^. Structural models of the dimeric proteins were predicted using AlphaFold3^15^.

### Cloning, heterologous expression and purification of proteins

*Ca*Kip3-NE-PT, PT, DT genes were PCR amplified from the Kip3N972_pET24d(+) plasmid^24^ and subsequently cloned into HT29 vector by Gibson assembly. HT29 is a pET16b-derived expression vector that adds 6xHis tag and maltose binding protein (MBP) fusion tags to the N-terminus of the expressed protein^49^. All plasmids were sequenced to verify correct construct assembly and gene sequences. *Ca*Kip3 motor domain genes were cloned previously in our laboratory^24^.

Plasmids were transformed into *E. coli* BL21 (DE3) cells, which were then grown in Luria-Bertani (LB) media supplemented with Ampicillin. Protein expression in these cells was induced with 1 mM isopropyl β-D-1-thiogalactopyranoside (IPTG) and grown overnight at 25°C in baffled flasks. Cell pellets were harvested using centrifugation and lysed by sonication in lysis buffer containing 50 mM sodium phosphate (pH 8), 500 mM NaCl, 5 mM 2-Mercaptoethanol, 0.2 mg/mL lysozyme, and Pierce Protease Inhibitor Tablets. Cell lysates were clarified by centrifugation at 35,000 x g for 25 minutes, after which the supernatant was loaded onto 10 mL of Ni-NTA resin (Qiagen) equilibrated with wash buffer (10 mM sodium phosphate, 300 mM NaCl, 20 mM imidazole, 5 mM 2-ME, pH 8). The resin was washed with 200 mL of wash buffer. The target protein was then eluted with elution buffer (wash buffer + 300 mM imidazole). Fractions containing the kinesin protein were identified by SDS-PAGE. Protein containing fractions were pooled, dialyzed overnight into HEPES buffer (20 mM HEPES, 150 mM NaCl, 1 mM dithiothreitol, pH 7.2), and loaded onto a Superdex 200 26/60 size-exclusion column (GE Healthcare) equilibrated with HEPES buffer. Fractions containing target protein were pooled, concentrated to 25 mg/mL with Amicon, flash frozen in liquid N_2_, and stored at −80 °C until use.

### Crystallization, data collection, and structure determination

Crystals of the *Ca*Kip3 PT construct that were suitable for X-ray diffraction data collection grew in 7 days from 5 μL hanging drops containing the protein (20 mg/mL) in a 1:1 ratio with a precipitant solution 0.1 M Tris pH 8.5, 18% polyethylene glycol (PEG) 6000, at 293 K. Prior to diffraction data collection, crystals were flash cooled in liquid nitrogen. Diffraction data were collected from a single crystal at beamline 08ID-1 of the Canadian Light Source (Saskatoon, Canada) at 100 K, and was indexed, integrated, and scaled with autoXDS. The Kip3 proximal tail structure was solved by molecular replacement using the AlphaFold3 predicted model of the *Ca*Kip3 PT monomer using MolRep^50^. The structures were refined with Refmac^51^ and Phenix Refine^52^ and manually optimized using Coot^53^. Data processing and refinement statistics are summarized in **Table 1**. Coordinates and structure factors of the *Ca*Kip3 PT structure have been deposited in the Protein Data Bank with the Accession Code 9CRW.

**Table 1.**
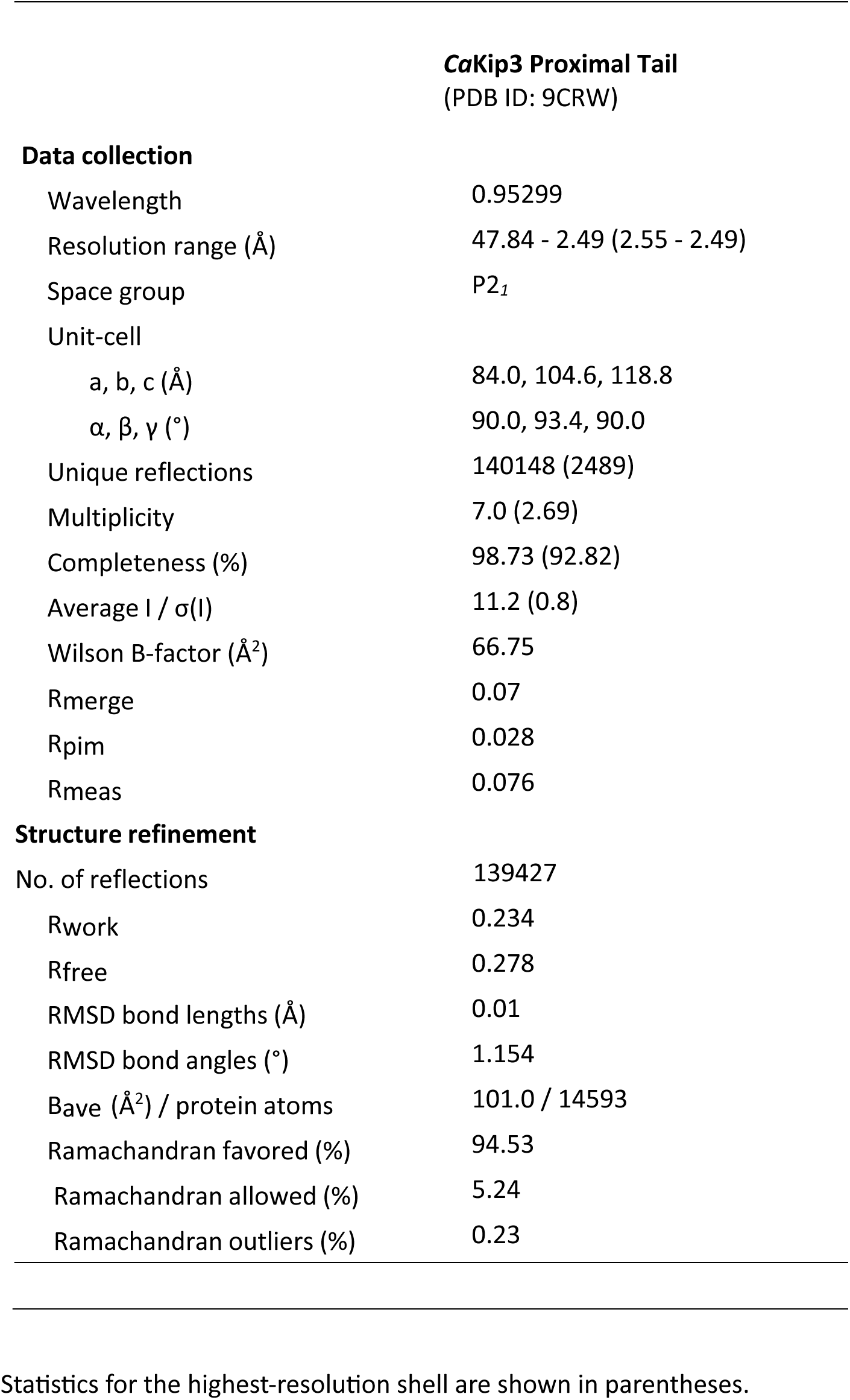
X-ray diffraction data collection, refinement, and validation statistics.

### Size-exclusion chromatography coupled with multi-angle light scattering (SEC-MALS)

Protein samples (2 mg/mL in PBS) were analyzed with static light scattering by injecting them into an AKTA Pure FPLC system with a Superdex 200 10/300 column (Cytiva). The chromatography system was coupled with a 3-angle light-scattering detector (TREOS II, Wyatt Technology) and a differential refractive index detector (OptilabT-rEX, Wyatt Technology). Masses (molecular weights) were calculated with ASTRA (Wyatt Technology).

### Analytical ultracentrifugation

Aliquots containing 20 μM protein in HEPES 20 mM, MgCl_2_ 1 mM, NaCl 150 mM, DTT 1 mM, pH 7.0 buffer were analysed in an XL-I analytical ultracentrifuge (Beckman Coulter, Fullerton, CA) equipped with an AnTi60 rotor. All samples were mixed thoroughly and allowed to equilibrate to 20 °C for 4 h before the start of the sedimentation analysis. 310 μL of samples were added to double-sector epon-filled centrepieces, with buffer alone in the reference compartments. Radial absorbance data were acquired at a wavelength of 290 nm, radial increments of 0.003 cm in continuous scanning mode, and a rotor speed of 40,000 rpm. Scans were taken at two-minute intervals. The resulting sedimentation boundaries were fitted to a continuous size distribution model (c(s)) using SEDFIT^54^. Size distributions were calculated using a radial grid of 500 values, a confidence level of p = 0.95, and fitted frictional ratio values.

### Microtubule co-sedimentation assay

The interaction between Kip3 tail domain constructs and microtubules was investigated with a robust kinesin microtubule equilibrium binding assay. In our experimental procedure, we first prepared 20 µM Taxol stabilized microtubules in BRB80 (80 mM PIPES, 1 mM MgCl_2_, 1 mM K-EGTA, 1 mM DTT, pH 6.8) buffer^24^. We pre-spun kinesin proteins at 155,000 x g for 25 min at 25 °C and used supernatant for further experiments Co-sedimentation reactions were prepared in 100-μL volumes of BRB80 buffer containing 2 mM adenylyl-imidodiphosphate (AMP-PNP), 1 μM kinesin, 20 μM taxol, KCl 100 mM, and varied concentration (0-10 µM) of microtubules. Reaction mixtures were incubated for 20 minutes at 25 °C then centrifuged at 155,000 x g (TLA-100; Beckman Coulter) for 25 minutes at 25 °C. An aliquot of the top supernatant fraction (50 μL) was collected, and the remaining 50 μL was discarded. Pellets were resuspended in 100 μL BRB80 buffer supplemented with 5 mM CaCl_2_, and 50 μL of the resuspension mixture was collected for SDS-PAGE analysis. Supernatant and pellet samples were mixed with an equal volume of 2x SDS loading buffer and 15 μL of each sample was run on a 10% SDS-PAGE gel and then visualized by staining with Coomassie brilliant blue R-250. Band intensities were quantified with ImageJ. The percentage of unbound versus bound kinesin was plotted as a function of microtubule concentration.

### SDS-PAGE and Western Blot Analysis of the *Ca*Kip3-PT construct

Crystals of the *Ca*Kip3 PT construct were harvested from a crystallization drop and dissolved in water. The protein solution was mixed with 2xSDS-loading buffer and subjected to SDS–PAGE. Gels were stained with Coomassie Blue dye or probed in Western Blot. Semi-dry Turbo transfer was used to transfer proteins on PVD membrane along with protein standards. Membrane was blocked with 1% milk solution and probed with Anti-6 x His Primary mouse antibody (Millipore #05-949, 1:1000 dilution) overnight at +4°C. Anti-Mouse DyLigh 680 secondary antibody from Invitrogen #35518 was used with 1:5000 dilution for 1h, RT. Detection was obtained at Odyssey DLx visualizing fluorescent signal of secondary Ab.

## Acknowledgements

This work was supported by funding from a Project Scheme grant from the Canadian Institutes of Health Research (PJT-169149) and an Individual Discovery grant from the National Sciences and Engineering Council of Canada grant (RGPIN-2019-05924).

## Author Contributions Statement

D.T., C.D., B.H., J.S.A, E.D. E.W and J.S.A modelled and analyzed the *Ca*Kip3 structure using AlphaFold and examined the sequence of *Ca*Kip3 using various bioinformatic tools; B.H., C.D. and D.T. cloned, expressed, and purified the kinesin proteins; D.T., J.S.A, E.D. and J.S.A collected X-ray diffraction data for the *Ca*Kip3 PT crystals, determined the structure, built and refined the atomic model, and analyzed the structure. D.T. performed the SEC-MALS analyses; D.T., C.D. and K.M. performed AUC analysis; D.T. and C.D. performed all other biochemical experiments; J.S.A. supervised the project; D.T., B.H., C.D. and J.S.A wrote the manuscript.

## Supplementary information

**Supplementary Figure 1.**
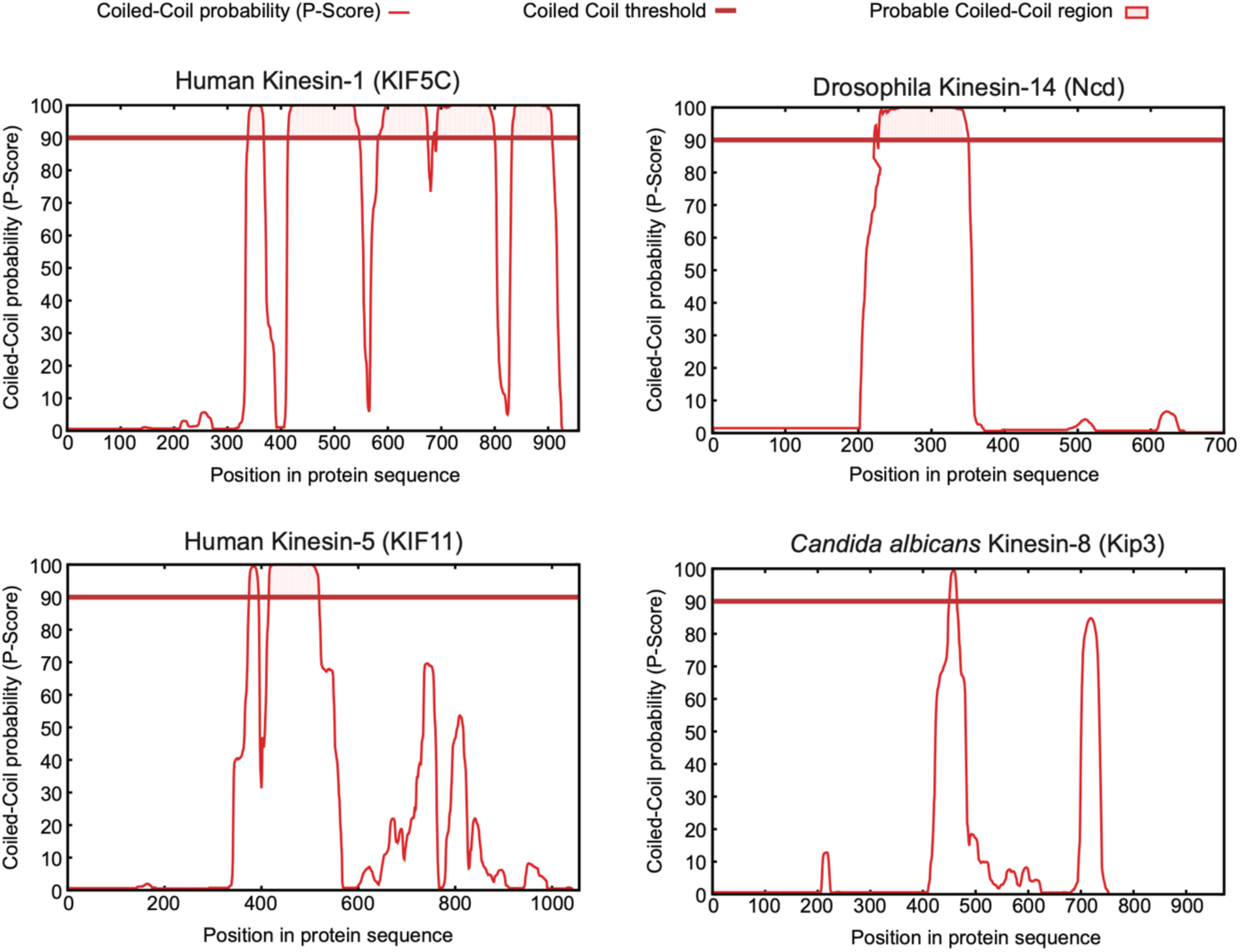
Coiled coil-forming propensity analysis of kinesins. The Waggawagga web-based tool for the comparative visualization of coiled coil predictions was used to illustrate the predicted coiled coil forming regions of the selected kinesin proteins^56^.

**Supplementary Figure 2.**
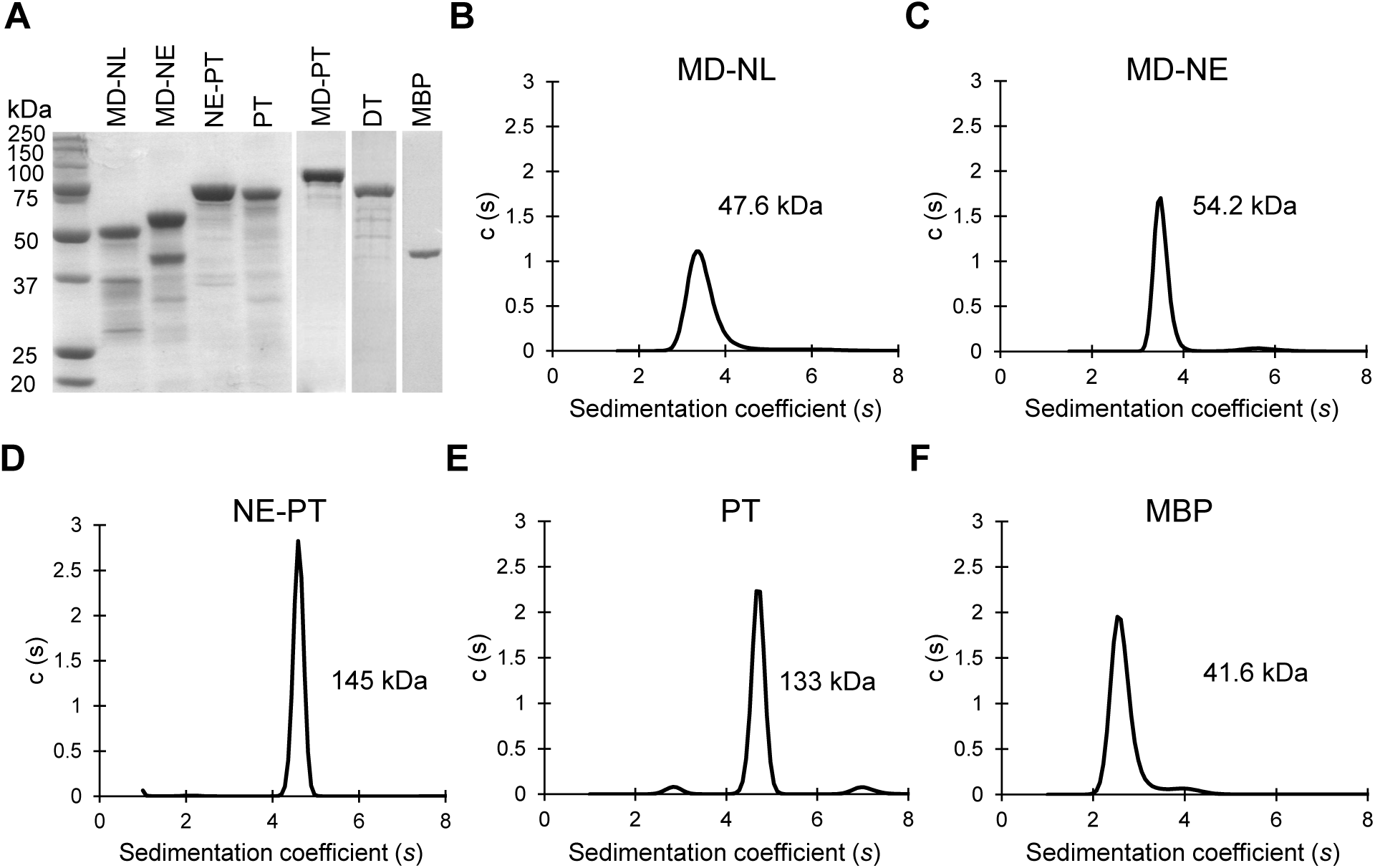
AUC analysis of the oligomeric state of *Ca*Kip3 constructs. (A) SDS-PAGE gel of all constructs stained with Coomassie dye. (B-F) Normalized sedimentation coefficient distributions c(s) determined by analytical ultracentrifugation (AUC) analysis of the 20 μM solution of Motor Domain+ Neck Linker (MD-NL), Motor domain + Neck Extension (MD-NE), Neck Extension + Proximal Tail (NE-PT), Proximal Tail (PT), and Maltose Binding Protein (MBP) in 20 mM HEPES pH 7.0, 1 mM MgCl_2_, 150 mM NaCl, and 1 mM DTT.

**Supplementary Figure 3.**
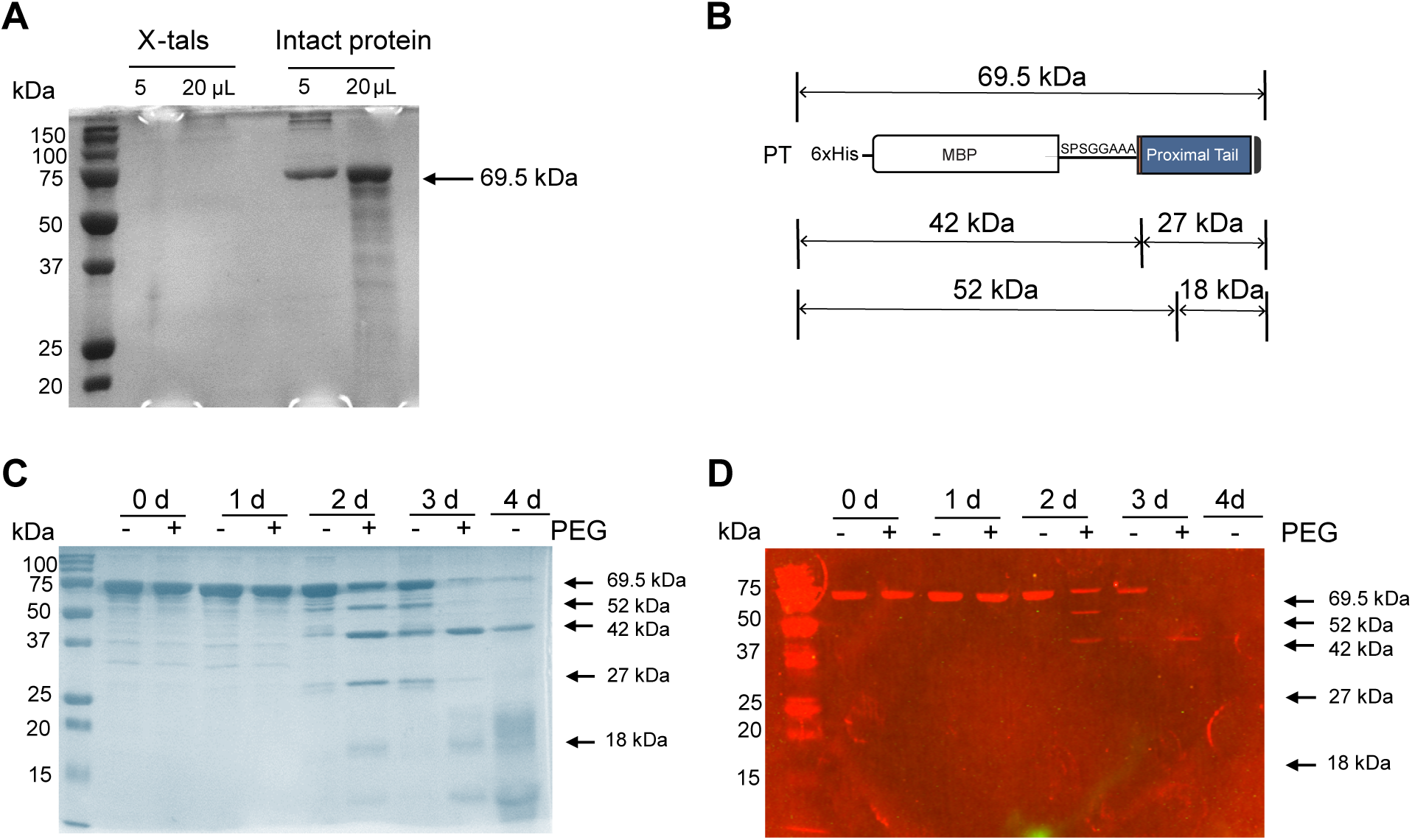
Analysis of the protein content of *Ca*Kip3 PT crystals. (A) Twenty-five crystals were harvested from crystallization drops with a loop, transferred to fresh well solution and then dissolved in water droplet containing 2xSDS loading buffer. The solution was transferred to an Eppendorf tube, heated at 80°C for 10 min, vortexed, and then 5 µl and 20 µl of it was loaded on 12% SDS gel. A control of intact protein was loaded alongside on the gel. The gel was stained with Coomassie dye, destained, and documented. (C) A 30 mg/mL solution of *Ca*Kip3 PT protein was mixed with 0.05 M Tris and 0 or 20% PEG 6000, pH 7 (to mimic crystallization droplet conditions (with PEG)) and control (without PEG). Both solutions were left at room temperature and 10 µL aliquots were removed after 1, 2, 3, and 4 days of incubation, diluted 30 times with water and mixed with 2xSDS loading buffer. Samples were analysed on 12% SDS gels and stained with Coomassie dye. (D) Western Blot of samples shown in (C). Semi-dry Turbo transfer was used to transfer proteins onto a PVD membrane along with protein standards. The membrane was blocked with 1% milk solution and probed with anti-6x His Primary mouse antibody (Millipore #05-949, 1:1000 dilution) overnight at +4C. Anti-Mouse DyLigh 680 secondary antibody from Invitrogen #35518 was used with 1:5000 dilution for 1h, RT. Detection was obtained at Odyssey DLx visualizing fluorescent signal of secondary Ab. (B) Diagram of potential proteolysis sites based on protein fragments detected in (C) and (D).

**Supplementary Figure 4.**
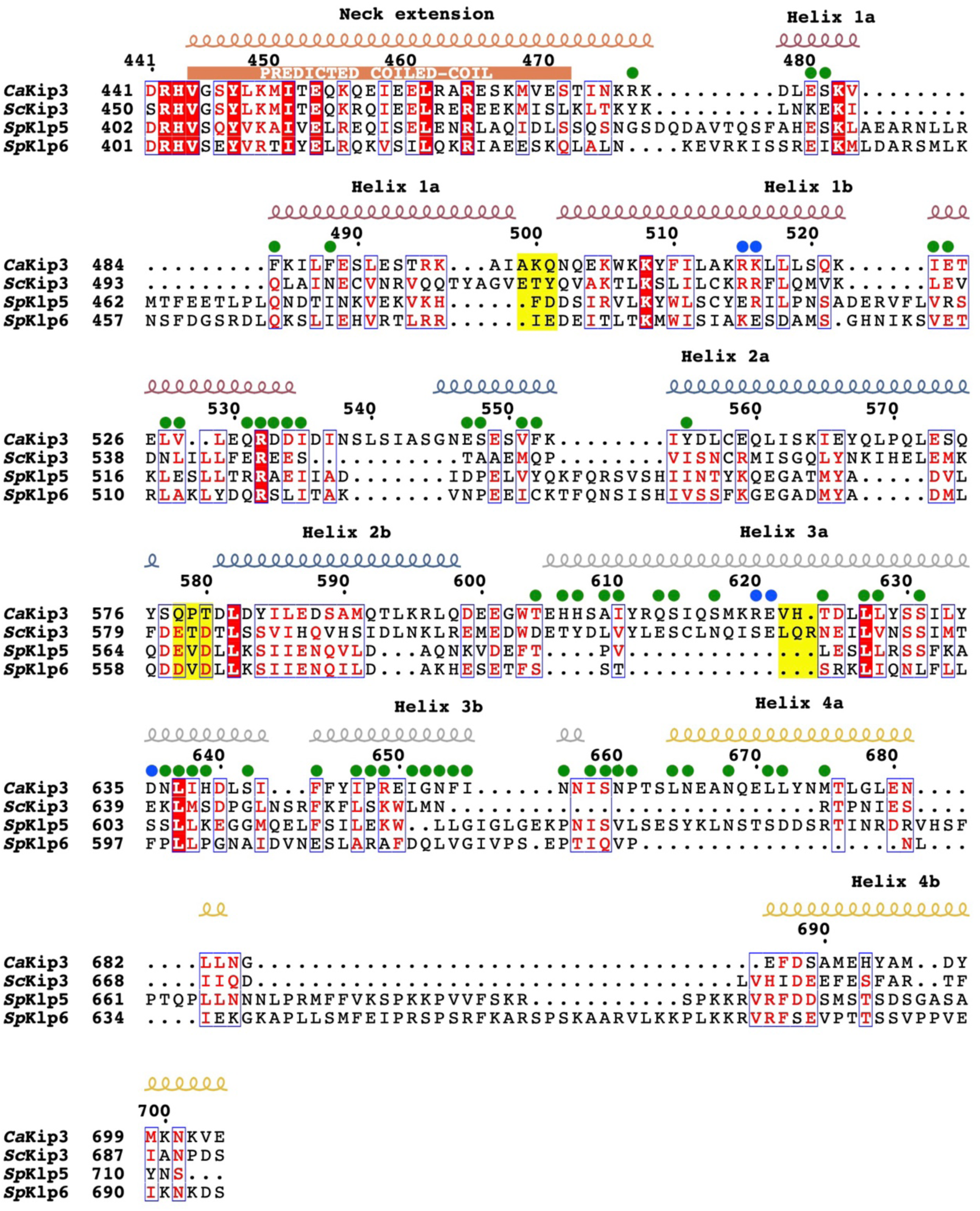
Sequence alignment of the neck extension and proximal tail of fungal kinesin-8s. The sequence alignment was performed using Clustl Omega^57^. Secondary-structure elements are placed above the sequence alignment according to the crystal structure of *Ca*Kip3 PT using ESPript^58^. Green dots are placed above amino acids that form intermolecular H-bonding, hydrophobic, or van der Waals interactions in the *Ca*Kip3 PT dimer. Blue dots are placed above amino acids that form salt bridging interactions in the *Ca*Kip3 PT dimer. Highlighted in yellow are regions that form the hinge domain of the helical bundle.

**Supplementary Figure 5.**
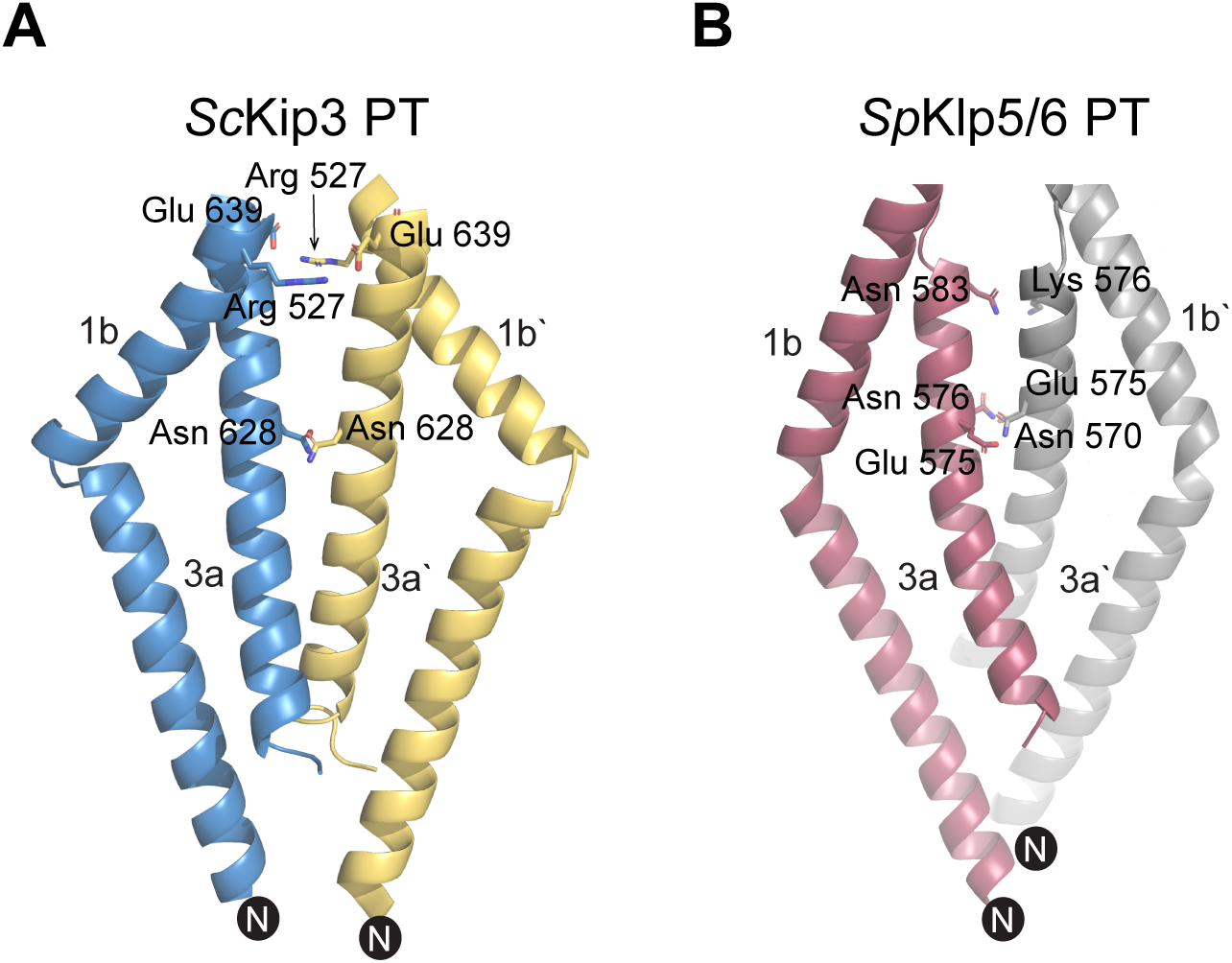
Salt bridge and polar interactions that stabilize the *Sc*Kip3 and *Sp*Klp5/6 proximal tail dimers. Cartoon representations of the main dimerization interface helices are shown for each AlphaFold3 model of the protein. Residues that form salt bridges or polar interactions are shown in stick representation and labelled.

